# Extrastriate activity reflects the absence of local retinal input

**DOI:** 10.1101/2023.05.15.540895

**Authors:** Poutasi W. B. Urale, Lydia Zhu, Roberta Gough, Derek Arnold, Dietrich Samuel Schwarzkopf

**Affiliations:** School of Optometry & Vision Science, University of Auckland, New Zealand; School of Psychology, University of Queensland, Brisbane, Australia; Queensland Brain Institute, University of Queensland, Brisbane, Australia

## Abstract

The physiological blind spot corresponds to the optic disc where the retina contains no light-detecting photoreceptor cells. Our perception seemingly fills in this gap in input. Here we suggest that rather than an active process, such perceptual filling-in could instead be a consequence of the integration of visual inputs at higher stages of processing discounting the local absence of retinal input. Using functional brain imaging, we resolved the retinotopic representation of the physiological blind spot in early human visual cortex and measured responses while participants perceived filling-in. Responses in early visual areas simply reflected the absence of visual input. In contrast, higher extrastriate regions responded more to stimuli in the eye containing the blind spot than the fellow eye. However, this signature was independent of filling-in. We argue that these findings agree with philosophical accounts that posit that the concept of filling-in of absent retinal input is unnecessary.

## Introduction

In the optic disc of the retina of each eye, there are mostly blood vessels and axons with no light-detecting photoreceptor cells, producing a physiological scotoma referred to as the blind spot. In visual space the blind spot is located near the horizontal meridian ∼15<u>°</u> of visual angle from the fovea and subtends roughly 6<u>°</u> by 8<u>°</u> (Awater et al., 2005; Li et al., 2014; Tootell et al., 1998). Despite making up a considerable discontinuity in retinal input, we are almost always unaware of our blind spot, even during monocular viewing (Figure 1A). Our visual system seemingly uses the intact visual field to interpolate into the blind spot, a process known as perceptual filling-in (Ramachandran, 1992). It is generally observed when spatiotemporally coherent elements abut or surround the scotoma (Fiorani Jr et al., 1992), or when a stimulus enters in from one side (Maus & Nijhawan, 2008; Maus & Whitney, 2016). Understanding perceptual filling-in is important because similar mechanisms could be at work with pathological scotomas after retinal damage (Gerrits & Timmerman, 1969; Ramachandran & Gregory, 1991; Zur & Ullman, 2003), and they may contribute to spontaneous or experience-dependent recovery from vision loss after brain lesions (Ajina & Kennard, 2012; Saionz et al., 2021). However, the neural mechanisms underlying this process remain largely unknown.

**Figure 1.**
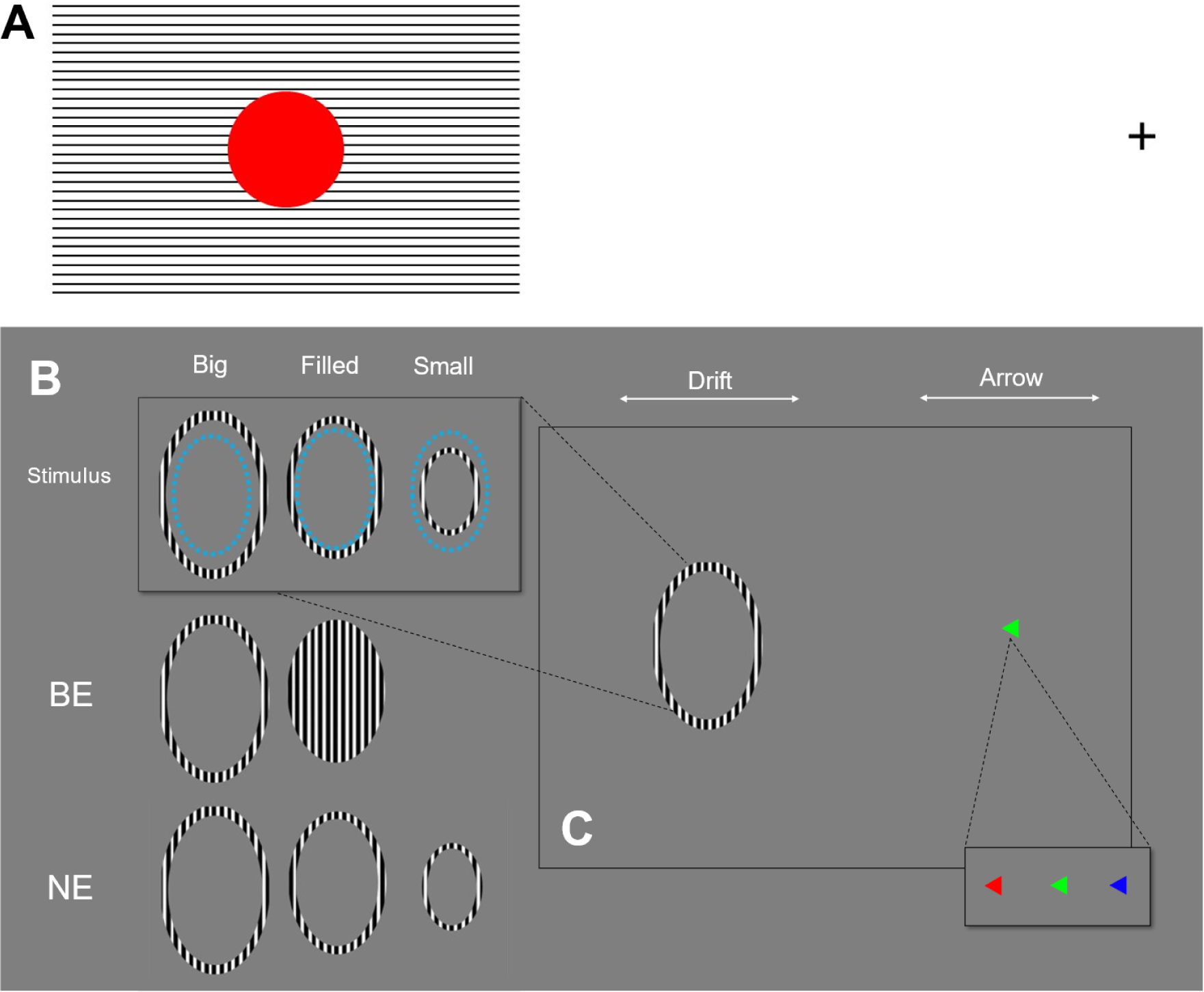
Demonstration of perceptual filling-in of the blind spot, and experimental stimuli and task. **A.** Demonstration of perceptual filling-in of the blind spot. If viewing on A4-sized page, this can be viewed by closing one’s right eye and fixating on the cross on the right at a viewing distance of 45cm. If the red circle is still visible, adjust the viewing distance (∼35-55cm) until the red circle is no longer visible, and the image on the left appears as a uniform striped grating. **B.** Stimulus conditions. Blue dotted outline shows blind spot boundary of hypothetical participant. Second and third rows show percept when viewing each condition through the blind spot eye (BE) and non-blind spot eye (NE), respectively. **C.** Layout of the screen as seen by a participant. The direction of the drifting sine-wave pattern and the color of the arrow, respectively, changed over the course of each experimental run.

Electrophysiological recordings of V1 neurons with receptive fields overlapping the blind spot have shown increased responses and tuning to line length suggestive of perceptually completed stimuli (Komatsu et al., 2000; Matsumoto & Komatsu, 2005). More recent research has shown that direction- and orientation-tuning remain in the V1 representation of the blind spot, irrespective of whether the incoming signal originate from the ipsilateral or contralateral eye (Azzi et al., 2015). However, the presence and magnitude of these signals does not match the intensity and completeness of the perceptual experience of filling-in. To our knowledge, no neuroimaging study to date has directly investigated potential neural correlates of perceptual filling-in of the physiological blind spot in human observers. Moreover, previous studies only focussed on the early visual cortex (Awater et al., 2005; Tong & Engel, 2001; Tootell et al., 1998), but the role of higher extrastriate regions in this process has never been tested.

Here we used our novel retinotopic mapping method (Urale et al., 2022) to identify activity within fMRI voxels encoding the blind spot location while stimulating the area within and outside of its boundaries, both during viewing with the blind spot eye (BE) and the non-blind spot eye (NE). We used a stimulus designed to elicit filling-in when viewing with the BE, but not the NE. Moreover, we used a control stimulus visible to both eyes and another that was only visible to the NE. Participants also performed a behavioural task to report whether they in fact experienced filling-in. A comparison of brain activity across conditions in both eyes and across stimulus conditions thus allowed us to investigate signals related to filling in: We hypothesized that visual brain regions responding more strongly when participants experienced filling-in should indicate a neural correlate of this illusory percept.

## Materials and Methods

### Participants

We recruited eight participants (ages: 21-43, 3 females, all right-handed) from staff and student populations. Each gave written informed consent to participate. Experimental procedures were approved by the University of Auckland Human Participants Ethics Committee (UAHPEC). The first session of this experiment involved data collected from seven participants for our work developing the reverse correlation pRF analysis (Urale et al., 2022), plus one additional participant.

### Procedure

Participants completed two scanning sessions: In Session 1 we conducted an experiment to retinotopically map the representation of the physiological blind spot and its surrounding region in human visual cortex. The results of this session and a comparison with conventional forward modelling pRF analysis are described in detail elsewhere (Urale et al., 2022). In Session 2 we measured Blood Oxygen Level Dependent (BOLD) responses with magnetic resonance imaging (MRI) while stimulating the area in and around the blind spot. In both sessions participants wore an ophthalmic eye patch over one eye (4 participants: left eye, 4 participants, right eye) prior to being placed in the MRI scanner bore. The other open eye would be the participant’s blind spot eye (BE). An MRI-compatible response pad with buttons was also placed into their dominant hand. Once in the bore, each session began with a behavioral localizer procedure (see below) that determined the dimensions and location of their physiological blind spot.

In both sessions, the localizer procedure was followed by the fMRI experiments. Once complete, we moved the patient bed slightly out of the bore and moved the eye-patch to the other eye, before conducting the same fMRI experiment but now with their control eye (NE). All stimuli were generated and displayed in MATLAB using Psychtoolbox 3 (Brainard, 1997; Pelli, 1997). Following this, we removed the participants from the bore and, in Session 1 only, attached a 32-channel head coil before replacing them into the bore and acquiring a T1-weighted structural scan of their brain anatomy.

### Experimental setup

Stimuli were presented on a liquid crystal display screen (BOLD screen, Cambridge Research Systems, Rochester, U.K.) placed at the rear of the scanner bore. Participants were able to view the screen via a mirror attached to the head coil at a viewing distance of 111cm. In degrees of visual angle, the screen subtended 35.5° vertically and 19.9° horizontally.

### Blind spot localizer

To determine the location and dimension of the stimuli during the retinotopic mapping and filling-in procedures, we first determined the location of the physiological blind spot in visual space. Once inside the scanner bore, participants fixated on a 0.17°×0.17° black dot on a halftone grey background (35.1 cd/m^2^) located 10.6° from the screen center on the side ipsilateral to their patched eye. To encourage participants to maintain fixation, the dot was framed by a radar grid pattern consisting of radial lines and concentric circles (Morgan & Schwarzkopf, 2019; van Dijk et al., 2016). The experimenter then used the computer mouse to move a red disc (diameter: 0.53°) across the screen, moving from within the blind spot to outside of the perimeter and vice versa. The participant was instructed to verbally communicate when they saw the disc reappear or disappear, at which point the experimenter would mark the current position of the red disc with a mouse click. The program then displayed a small grey dot at that location. This was repeated until a clear shape was defined by the grey dots. If satisfied, the experimenter pressed the right mouse button, prompting the program to use the coordinates of the grey dots to calculate the centroid of the estimated blind spot location, displaying a black dot (diameter: 0.17°), nested inside the outline of a larger circle (diameter: 10.6°). For Session 1, this area would be stimulated in the forthcoming pRF mapping fMRI runs. For Session 2, these same coordinates were used to generate an area that matched the shape of the blind spot during the filling-in experiment.

### Session 1: Retinotopic mapping of the blind spot

For retinotopic mapping in Session 1, we used similar stimuli to previous studies using pRF mapping (Alvarez et al., 2015; Dumoulin & Wandell, 2008; Morgan & Schwarzkopf, 2019; Schwarzkopf et al., 2014), with some key differences. Participants fixated on a dot located 11.5° from the center of the screen on the side ipsilateral to the NE eye. The dot was framed by the same radar grid pattern from the blind spot localizer on one side of the screen. Bar stimuli (width: 1.3°) traversed a circular centered on the blind spot, as defined by the blind spot localizer. This bar contained a checkerboard pattern (side length: 0.95°) that flashed at 6Hz. The bar traversed the circular region in eight orientation/directions, beginning in a horizontal orientation and moving upward (0°), and then changing in increments of 45° until an orientation of 315°. The bar moved at a rate of 0.4° per second and took 25 seconds to traverse the circular region in one direction. Because the bar appeared within a circular aperture, its length changed during each sweep. Between the fourth and fifth sweep and at the end of the eighth sweep, we presented a blank period where only the fixation dot and radar pattern were visible.

To ensure that participants fixated on the black dot, we required them to perform a detection task. Every 200 ms, there was a 10% probability that the fixation dot would be replaced with a number or letter (letters A-Z and numbers 0-9) for a duration of 200 ms. Participants were asked to press a button on the response pad when they saw a number. The 200 ms epoch immediately following the presentation of a number always showed a fixation dot. We collected the same number of retinotopic mapping runs for each eye (see below).

### Session 2: Re-localization of blind spot and filling-in experiment

In Session 2 participants again underwent the blind spot localizer procedure before beginning the filling-in experiment. The aim of this experiment was to record BOLD responses while stimulating the area in and around the blind spot during viewing through the BE and NE. To stimulate the blind spot, we generated several ring stimuli (Figure 1B).

### Determining stimulus conditions

Before the filling-in experiment, we first conducted a procedure to determine the stimulus properties for three conditions: A “big” condition, consisting of a stimulus equally visible to both the BE and NE, a “filled” condition abutting the blind spot’s boundary and thus elicit filling-in only in the BE, and a “small” condition located inside of the blind spot and therefore only visible in the NE.

The dimensions of these conditions were determined inside the scanner. Participants were fitted with an eye-patch over their NE and presented with an irregularly shaped ring stimulus matching the blind spot boundary estimated in Session 2. The ring was 0.9° thick with equal thickness falling within and outside the blind spot boundary. Its boundary was hard-edged and filled with a vertical sine grating (spatial frequency = 3.45 cycles/degree, temporal frequency = 7 Hz, maximum luminance: 68.5 cd/m^2^; minimum: 0.1 cd/m^2^) that moved either left or right (Figure 1B). Participants fixated on an arrow at the same location as the fixation dot from the localizer procedure while the ring stimulus was presented to the area corresponding to their blind spot. Participants were asked to describe whether the ring was visible at all, and if so, whether they were experiencing the stimulus as a continuous patch (i.e., filling-in), as an outline, or if only part of the ring was visible (i.e., due to the ring’s boundary falling partially within the blind spot). In verbal coordination with the participant, the experimenter used a mouse and keyboard to adjust the scale and location of the ring stimulus until participants reported seeing filling-in. These stimulus dimensions were stored and comprised the ring dimensions in the “filled” condition. Participants were asked to remember the sensation of filling-in for forthcoming tasks. Following this, the scale of the stimulus was increased to the point that participants reported only seeing a ring, and not a complete patch (“big” condition), and subsequently decreased until it was no longer visible (“small” condition). Finally, the participant was shown each condition again and reported what they saw. If prompted, the experimenter communicated with the participant until the desired perceptual effect of each condition was achieved. Once satisfied with each condition, participants remained in the scanner for the filling-in experiment.

### Filling-in experiment and behavioral task

For the filling-in experiment participants completed a series of 300-second runs. Each run consisted of successive presentations of the three ring stimulus conditions plus blank intervals. Each condition was presented for 15 seconds. The three ring conditions occurred in successive pseudo-randomized triplets (e.g., filled, small, big) separated by a blank interval. Each condition occurred five times per run. Each run was divided into epochs of 3 seconds. At the beginning/end of each epoch occurring during presentation of a ring stimulus, the direction of the drifting grating changed (left or right). Concurrently, the fixation arrow had a 50% chance of switching directions from left to right (or vice versa). To avoid adaptation, the arrow cycled through three colors (red, green, blue) at the start of each epoch.

During each run, participants were asked to perform tasks designed to encourage fixation to the arrow and attention towards the ring stimulus (when present). They were instructed to press a button (positioned at their index finger) once if they saw the fixation arrow change direction, and to press twice in close succession (“double press”) if when the arrow pointed in the same direction as the movement inside of the ring stimulus. For example, participants were to press the button once if the arrow changed from left to right while the drifting pattern was perceived as moving leftward (or if it was not visible at all) or press twice if the drifting pattern moved rightward. Thus, single presses represent fixation compliance (*arrow subtask*), and double presses imply attention encompassing both the fixation arrow and the ring stimulus. Concurrently, participants were instructed to press a second button (positioned at their middle finger) once if they experienced filling-in (*filled task*).

Following several runs (3-5) while viewing through the BE, participants were partially removed from the scanner bore to shift the eye patch to the other eye, before completing the same number of runs viewing with the NE. Instructions in were identical during runs while viewing through the BE and NE.

### Magnetic resonance imaging

We used a Siemens MAGNETOM Skyra 3 Tesla scanner with a 32-channel head coil based at the Centre of Advanced Magnetic Resonance Imaging at the University of Auckland. During all functional imaging, we removed the front element of the coil to allow for an unrestricted field of view. The setup featured 20 effective channels covering the back and sides of the head. In Session 1, we performed 4-6 pRF mapping runs of 250 T2*-weighted image volumes in each eye and an accelerated multiband sequence with a TR of 1000 ms, 2.3 mm isotropic voxel resolution, and 36 transverse slices. These slices were angled to be roughly parallel with the calcarine sulcus. The scan had a TE of 30 ms, a flip angle of 62°, a 96 × 96 field of view, a multiband/slice acceleration factor of 3, and in-plane/parallel imaging acceleration factor of 2, and rBW of 1680 Hz/Px. Following the pRF runs, the front element of the coil was attached to achieve optimal signal-to-noise levels for collecting a structural scan. This scan was a T1-weighted full-brain anatomical magnetization-prepared rapid acquisition with a gradient echo (MPRAGE) scan with a 1 mm isotropic voxel size. The scan procedure in Session 2 was identical to Session 1, except with 3-5 runs per eye (i.e., 6-10 runs total) each with 300 T2*-weighted images.

### Pre-processing

For both sessions we realigned and co-registered functional data to the anatomical scan in SPM12 (Wellcome Centre for Human Neuroimaging; https://www.fil.ion.ucl.ac.uk/spm) using default parameters. Following this, we used the automatic reconstruction algorithm on the structural scan in FreeSurfer (Version 7.1.1: https://surfer.nmr.mgh.harvard.edu) to reconstruct inflated surface mesh models of the boundary between grey and white matter and the boundary between grey matter and the pia mater (Dale et al., 1999; Fischl, 2012; Fischl et al., 1999). We then projected functional data onto this model by locating the voxel that sat on the mid-way point between each vertex on the grey-white matter boundary and the same vertex on the pial surface boundary. To remove slow drifts, we applied linear detrending to the time series for each vertex and run, and then normalized the time series to z-scores. We averaged runs from the BE and NE, respectively, leaving a single run for each eye with 250 volumes each. For the Session 2 data, we instead concatenated the time series for all runs across both eyes.

For Session 1 data, we limited further analyses (see pRF analysis below) to an area approximately located in the occipital lobe by selecting vertices in the inflated cortex model with FreeSurfer y-coordinates of ≤ -35. Next, we determined a noise ceiling to identify vertices with a reliable response to visual stimuli, by correlating the time series between even and odd runs in each vertex, separately for the BE and NE conditions. We used this split-half correlation (*r*) and the Spearman-Brown prophecy formula (Spearman, 1910) to calculate a noise ceiling:

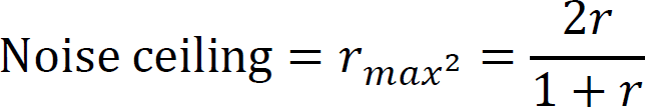

The noise ceiling therefore represents the maximum R^2^ achievable for a given time series, and shows the extent that a given vertex is visually responsive. To select only vertices with a reliable visual response, subsequent pRF analyses were limited to vertices with a noise ceiling >.15 (see example maps in Figure 2A). As expected, this revealed small clusters of visually evoked responses in the early visual cortex because we only stimulated a small region of the peripheral visual field.

**Figure 2.**
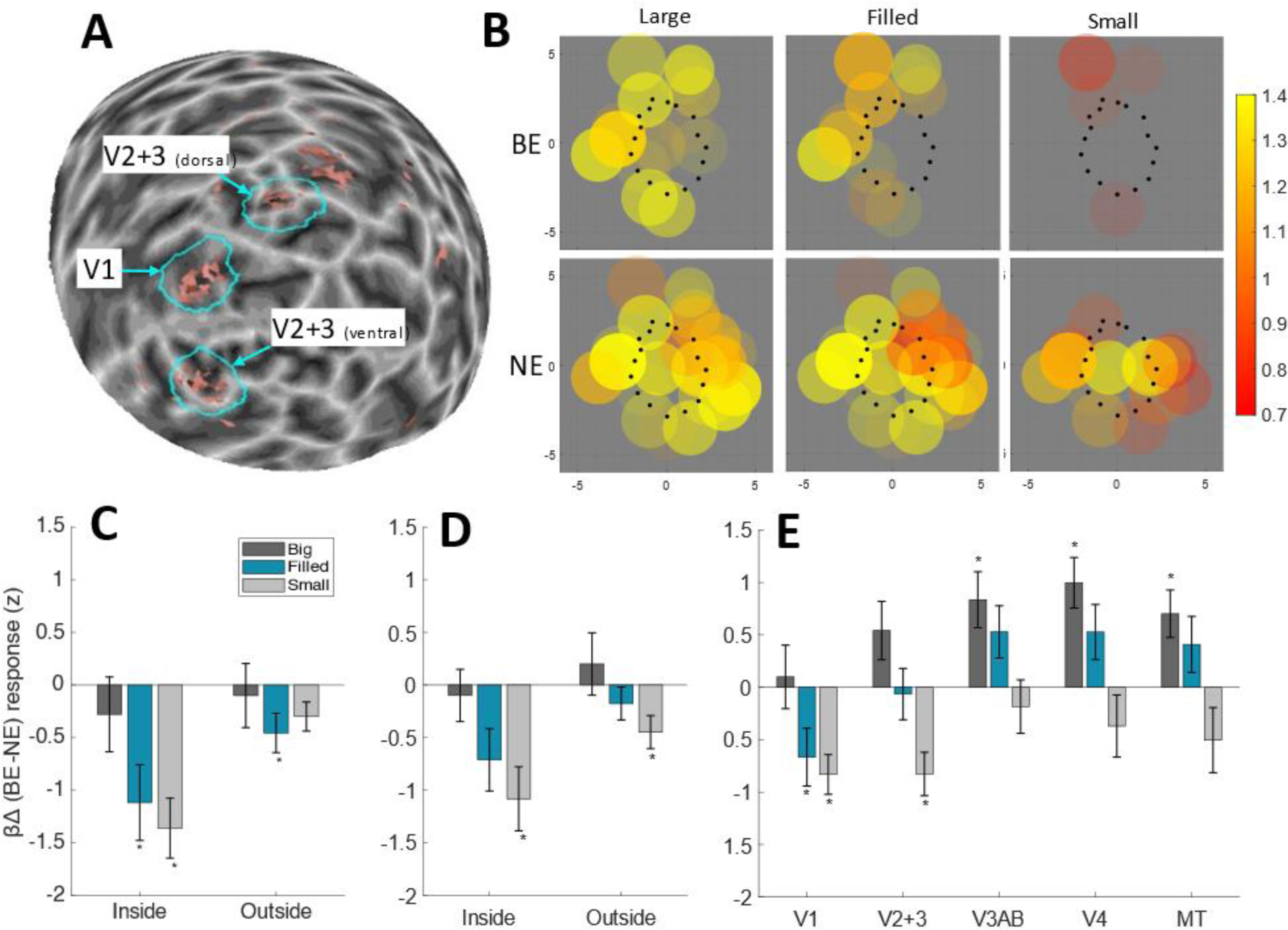
Neural responses to a filling-in stimulus. **A.** Retinotopic areas showing robust visual responses during retinotopic mapping (noise ceiling >0.15) shown on spherical reconstruction of the grey-white matter boundary of the right occipital lobe of one participant. Shades of grey indicate gyri and sulci. The light blue lines indicate regions of interest determined by the geodesic selection procedure (see Methods for details). **B.** Reconstruction of retinotopic V1 responses to each experimental condition in an example participant (N99). Circles show location, size, and response amplitude (see color code in arbitrary units) of individual pRFs in and around the blind spot during Session 2 as determined by the retinotopic map in the control eye as mapped in Session 1 (i.e., V1_pRF_). Only pRF responses (β) >.7 are shown for clarity. Coordinates are in degrees of visual angle relative to the blind spot center. The top and bottom rows show data from blind spot eye (BE) and non-blind spot eye (NE) viewing, respectively. Black dots show the location of the blind spot boundary from the behavioral localization procedure. **C-E.** Mean difference in response to BE and NE stimulation (BE-NE βΔ) for each stimulus condition within individual regions of interest. V1_pRF_ **(C)** and V2+3_pRF_ **(D)** data, separated into subregions inside and outside the blind spot. **E.** Mean difference in responses for regions of interest defined based on visual responsivity. Error bars represent ±1 standard error of the group mean. Bars labelled with a * differ significantly from zero (one-sample t-test against zero, false discovery rate-corrected p < .05).

### pRF analysis

We performed pRF analysis on Session 1 scan data for the purpose of identifying vertices encoding the area inside and around the blind spot. These procedures are identical to those used in our previous work (Urale et al., 2022). Population receptive field locations (x, y) and size (σ) were modelled using SamSrf Toolbox (Schwarzkopf, 2018). We treated the center of the blind spot as the center of the (mapped) visual field and used moving bar stimuli to stimulate a circular area with a radius of 5.3° around it. We created multiple stimulus apertures indicating the location of the bar stimulus during the recording of each 1 s fMRI volume, and generated 100×100 pixel matrix where the activity in each pixel corresponded to the presence of a stimulus at the corresponding location in the visual field during the scan. We accounted for the lag in blood oxygenation-dependent response by convolving this aperture with a canonical hemodynamic response function from previously collected data (de Haas et al., 2014).

We then regressed the time series of each vertex on the time series of each pixel in this matrix. The result was 100×100 of β coefficients for the observed time series from each vertex. The mosaic of coefficients for each vertex therefore represented its responsiveness to areas across the stimulated visual field, i.e., its pRF profile. The maximum of the pRF profile was taken as the peak response pixel. Further analyses were limited to vertices for which the squared correlation between this pixel and fMRI response was greater than 0.1. Additionally, the size and spatial location of the profile were determined by fitting a symmetric two-dimensional Gaussian profile, with x and y coordinates, standard deviation, and amplitude (β_pRF_) as free parameters. Analyses were further restricted to vertices where goodness-of-fit of the Gaussian profile R^2^>0.5. This left vertices that were reliably responsive to the bar stimulus, and which were determined as having a sufficiently clear pRF profile. Moreover, to save computation time, the pRF analysis was limited to vertices in the occipital lobe and only to those vertices showing a robust visual response (see Pre-processing).

### General linear model

The time series of the filling-in stimuli from Session 2 were concatenated and included in a general linear model (GLM) using SamSrf. Boxcar regressors defined each condition (big, filled, and small for the BE and NE, respectively) in each run, and convolved with the canonical HRF (de Haas et al., 2014). Additionally, six motion regressors (x, y, z, pitch, roll, yaw) and a global constant for each run were included in the GLM. For all conditions, signal was calculated by summing the β weights for the presentation of the various ring stimuli and contrasting conditions against one another. We also calculated the overall response to the big ring condition irrespective of the eye-of-origin. We then smoothed these activation maps across the spherical surface mesh (full width at half maximum of 3 mm) for the purpose of selecting regions of interest.

### Regions of interest

We used two separate procedures to determine regions of interest (ROIs). To analyze the retinotopic representation of the blind spot and its surround, we used results of the pRF analysis from Session 1; for each participant, we projected pRFs from the NE condition limited to those with a good fit pRFs (R^2^ >.5) and manually selected the vertex falling at the center of the cluster in V1, and the ones in dorsal and ventral V2 and V3, respectively. A geodesic region growing procedure defining a boundary 12 steps from this center location across the grey-white matter surface mesh resulted in a roughly circular patch around each cluster. Because the blind spot intersects the horizontal meridian, corresponding to the border between V2 and V3, we only defined a combined ROI in those areas. These pRF-derived ROIs are denoted with the “pRF” subscript, i.e., V1_pRF_, V2+3_pRF_. Moreover, we used the NE retinotopic map together with a convex hull of the blind spot localizer data to specify which pRFs fell within and outside the blind spot. Note that in V2+3_pRF_ for participant N02, no pRFs were identified inside the blind spot and this participant was therefore excluded from analysis for this subregion of interest.

Following the retinotopic analysis, we conducted a coarser analysis of visually responsive brain regions irrespective of retinotopic specificity. ROIs were defined by first identifying areas responsive to the big ring stimuli, regardless of the eye-of-origin. We used smoothed maps of the overall response to the big stimulus for either eye in conjunction with brain areas defined by an atlas (Sereno et al., 2022) derived from phase-encoded retinotopic, tonotopic, and somatosensory, and motor fMRI experiments (Dick et al., 2012; Huang & Sereno, 2013, 2018; Sood & Sereno, 2016, 2018). The visual areas in this atlas are based on group average retinotopic mapping data that tested the visual field up to an eccentricity of 50° at all polar angles, as well as T1 mapping data, with parcellation based on topographic features first defined through manually defining cortical surfaces based on previously established boundaries from literature defining a total of 57 visual areas (for a full list, see (Sereno et al., 2022), Table 1). We then combined several of these individual atlas ROIs into larger clusters (Supplementary Table 1). Visual areas that showed consistent visual activation across all participants were selected for subsequent analyses. A total of five areas were selected: V1, V2, V3, V3A/B, V4, and MT. While responses were present in other areas, these were either inconsistent across participants or unlikely to signal visually evoked responses (Davey et al., 2016; Raichle, 2015). Within each of the selected visual areas in the atlas, a ROI was generated by selecting a contiguous cluster of vertices around the peak response using a region growing approach containing coefficients greater than the 98^th^ percentile of β values. These ROIs are referred to with the “GLM” subscript, i.e., V1_GLM_, V2+3_GLM_, V3A/B_GLM_, V4_GLM_, MT_GLM_.

## Results

We used functional magnetic resonance imaging (fMRI) combined with highly accurate retinotopic mapping (Urale et al., 2022) to investigate neural responses in the cortical representation of the physiological blind spot. We hypothesized that populations of visual neurons activated preferentially while participants experienced filling-in constitute a neural correlate of this percept. First, we behaviorally delineated the borders of the scotoma in individual participants (Supplementary Figure S1). Then we measured neural responses while participants viewed stimuli designed to elicit filling-in when viewing with the blind spot eye (BE), but not the non-blind spot eye (NE). Moreover, we used a big control stimulus equally visible to both eyes, and a small control stimulus only visible to the NE. Comparing brain activity across conditions in both eyes and across stimulus conditions thus allowed us to investigate signals related to filling-in. Participants also performed a behavioral task that indicated they reliably experienced filling-in for the medium sized stimulus (Figure 1B-C and Supplementary Information). Note, while some participants did not report filling-in during all such trials, the small number of trials without this percept precluded us from analyzing these trials separately.

We then tested the preferential response of neuronal populations in early visual cortex representing the area in and around the blind spot during viewing through the BE versus the NE (Figure 2A). Figure 2B shows activity measured for individual V1 population receptive fields (pRFs) of one participant, split by eye and stimulus condition. There were fewer responsive pRFs in any of the BE versus NE conditions. Importantly, pRFs whose centers fell inside the blind spot did not respond at robust levels during BE viewing at all. To quantify this across all participants, we performed a group analysis of stimulus conditions (big, filled, and small). For each pRF, we calculated a difference score indicating stronger activation to BE than NE stimulation (BE-NE; βΔ). We further divided pRFs into subregions depending on whether they were within or outside the blind spot. Then we averaged these values across all pRFs within each subregion (Figure 2C,D). Responses in both V1_pRF_ and V2+3_pRF_ were stronger in the NE than the BE for the filled and small conditions. In V1_pRF_, responses differed significantly with location relative to the blind spot boundary (repeated measures analysis of variance: F(1,7)=8.95, p=.02), and between the three stimulus conditions (F(2,14)=4.54, p=.03). The difference in responses to the three conditions also depended on location (interaction: F(2,14)=3.81, p=0.048). We observed similar results for V2+3_pRF_ (location effect: F(1,6)=12.75, p=0.012; condition: F(2,12)=4.6, p=.033; but no interaction: F(2,12)=0.63, p=0.55).

Crucially, pRFs inside the blind spot in V1_pRF_ responded significantly less to BE than NE stimulation in the filled and small conditions (one-sample t-tests vs. 0; small: t(7)=-4.77, p=0.002; filled: t(7)=-3.11, p=0.017; α=0.0292 corrected for false-discovery rate), but not the big condition (t(7)=-0.78, p=0.459). In V2+3_pRF_ this difference was only significant for the small condition (t(6)=-3.56, p=0.012) but not the other stimulus conditions (filled: t(6)=-2.4, p=0.053; big: t(6)=-0.4, p=0.703). The big stimulus was equally visible through both eyes, and therefore should result in similar responses in either eye. Due to its size, this stimulus likely drove neuronal populations outside the retinotopic region of interest, yet it still marginally activated the cortical representation of the blind spot. The rationale of our experiment was that these effects should be robustly identifiable at the level of individual participants. We therefore defined the minimum criterion for consistent effects that at least 6 of the 8 participants must show the same significant difference as the group average (one-sample t-test vs zero, p<.05, Bonferroni-corrected by the number of conditions and regions of interest). The filled and small stimulus condition met this criterion in V1_pRF_, while in V2+3_pRF_ this was the case only for the small condition. The cortical subregion outside the blind spot representation also responded either the same or only mildly less during stimulation of the BE, except for the filled stimulus in V1_pRF_ and the small stimulus in V2+3_pRF_ both of which responded significantly less to the BE. At the individual level, these same conditions also met our consistency criterion. Some small errors in pRF position estimates are inevitable, and some pRFs will also straddle the blind spot border. One should therefore expect small differences in activation.

Importantly, results from early retinotopic regions stand in stark contrast to activity measured in higher extrastriate areas selected for their general visual responsivity. Here we found stronger responses while stimulating the BE compared to the NE in both the filled and big conditions (Figure 2E). The broad retinotopic specificity in these areas meant we could not parse apart neuronal populations mapped to the inside and outside of the blind spot. For both the big and filled conditions, BE responses were greater than NE responses in V3A/B_GLM_, V4_GLM_, and MT_GLM_, although this difference was only significant for the big condition. Meanwhile, results for V1_GLM_ and V2+3_GLM_ were consistent with those from retinotopic regions of interest with pRFs inside the blind spot boundary. Specifically, responses differed between stimulus conditions (F(2,14)=15.89, p<0.001) and also between regions (F(4,28)=5.13, p=0.003), and there was an interaction between these effects (F(8,56)=2.28, p=0.034). Because one participant potentially failed to comply with task instructions, we repeated this analysis after removing their data, but found the overall pattern remained qualitatively the same (Supplementary Figure S2 and Supplementary Information). We also repeated our consistency analysis for these data. For all regions of interest, we found weaker responses to the small stimulus in the BE in at least 6 participants. In V1_GLM_, the filled stimulus also met this criterion. Meanwhile, the big stimulus produced consistently stronger BE responses in V4_GLM_ for all participants, and in MT_GLM_ both the big and filled stimuli produced stronger responses in at least 6 participants.

## Discussion

Overall, these results suggested no evidence of a neural correlate of an illusory percept in early retinotopic areas mapped to the location of the blind spot. Rather, these regions merely signaled the retinal input, not the perceived filled-in space. This disagrees with previous research suggesting early visual areas as a locus for filling-in (Fiorani Jr et al., 1992; Komatsu et al., 2002; Matsumoto & Komatsu, 2005; Ramachandran, 1992; Spillmann et al., 2006). Single-cell recordings in monkeys (Azzi et al., 2015; Komatsu, 2006; Komatsu et al., 2000, 2002; Matsumoto & Komatsu, 2005) have suggested that some V1 neurons show sustained spike rates while observing stimuli known to elicit filling-in in humans. It is possible that we could not detect such signals because they are weak or due to the paucity of such neurons. However, even such signals would not be at the level of activity expected by the filled-in percept of a high contrast grating. Our findings also demonstrate the accuracy of our retinotopic mapping procedure: pRFs identified as being inside versus outside the blind spot responded as predicted by their estimated visual field location.

In contrast to early regions, we found stronger visually evoked responses through the blind spot eye (BE) in several extrastriate regions beyond V2/V3. We posited such a response pattern as a signature of perceptual filling-in because it indicates the presence of a stimulus that was not there. However, our findings do not support the interpretation that these higher extrastriate responses are a neural correlate of filling-in. As hypothesized, a neural correlate should only appear for the filled condition, where the stimulus abutted the blind spot border and participants experienced a filled grating. The big condition was equally visible through both eyes and participants rarely reported filling-in (most likely erroneous button presses); however, we measured the same signature of stronger BE responses also for this condition, and in fact this was even greater than for the filled condition. What then could cause such pronounced differences in responses between the eyes?

One explanation is that the gap inside the ring stimulus modulates higher visual areas differently, depending on the eye. Filling-in could occur due to the absence of signals from retinotopic locations mapped to inside the blind spot. Combined with a signal outside its borders (as in the big and filled conditions) there is less modulation compared to NE viewing. Receptive fields in higher-level regions are large (Dumoulin & Wandell, 2008) and encompass the blind spot as well as its surrounding space. Greater activity in these regions could indicate that rather than an active filling-in process, higher extrastriate neurons simply discount the missing input, because the monocular neurons in the corresponding part of V1 do not respond. The process might also factor in the luminance difference between the inputs from the two eyes. Intrinsically photosensitive retinal ganglion cells containing melanopsin (Dacey et al., 2005; Gamlin et al., 2007; Lucas et al., 2003; Tu et al., 2006) could play a role, as they might mediate the pupil light reflex elicited by stimulation of the physiological blind spot (Miyamoto & Murakami, 2015; Saito et al., 2018). The detection of luminance stimulation could provide an error signal to extrastriate cortex to ignore the lack of patterned input.

The stronger differential responses we observed for the big condition likely reflects the greater stimulus energy in the big condition during BE-viewing, combined with the lack of inhibitory input from inside the blind spot. The filled condition during BE viewing is subject to the same absence of input, but the overall stimulus energy is lower; consequently, the net difference between BE and NE is smaller.

Our suggestion that higher regions discount the missing input bears similarities to earlier explanations of filling-in (Dennett, 1992, 2017; Durgin, 1998; Durgin et al., 1995) that in the absence of information, higher visual areas simply label the missing area as “more of the same”. Many cognitive scientists have rejected that idea in favor of filling-in as an active process; however, we argue that previous findings are in fact consistent with this explanation (Churchland & Ramachandran, 1996). Spatial integration by higher visual areas in the absence of local retinal input could operate at an intermediate stage, labelling local surfaces and simple contours, not more complex objects and textures. This would explain why the visual system perceptually completes a line that spans the blind spot, but not the curvature of a ring (Ramachandran, 1992), that it operates at the level of second-order contours (Figure 3A), and other similar findings (Crossland & Bex, 2009; Spillmann et al., 2006). One argument levelled against Dennett’s account, is that arrays of stimuli as in Figure 3B are not filled in globally (Churchland & Ramachandran, 1996). However, this is precisely what a discounting process at intermediate stages of visual processing predicts: Our perception interprets the donut hole falling in the scotoma as a complete disc, but it does not go so far as to replace the percept with distal information of more donuts. Interestingly, however, the uniformity illusion (or “healing grid”), an effect that does not rely on V1 (Suárez-Pinilla et al., 2018), does even out inhomogeneities in peripheral stimulus arrays in our perception (Otten et al., 2017). This process could also play a role in filling-in of the blind spot under some circumstances. On the other hand, electrophysiological studies reported signals of V1 neurons with receptive fields overlapping the blind spot suggestive of perceptually completed stimuli (Komatsu et al., 2000; Matsumoto & Komatsu, 2005); these signals could however be an epiphenomenon of this discounting process and/or result from feedback connections. In any case, such signals were generally found to be subtle, not at the magnitude the experience of the actual perceived filled stimulus would predict.

**Figure 3.**
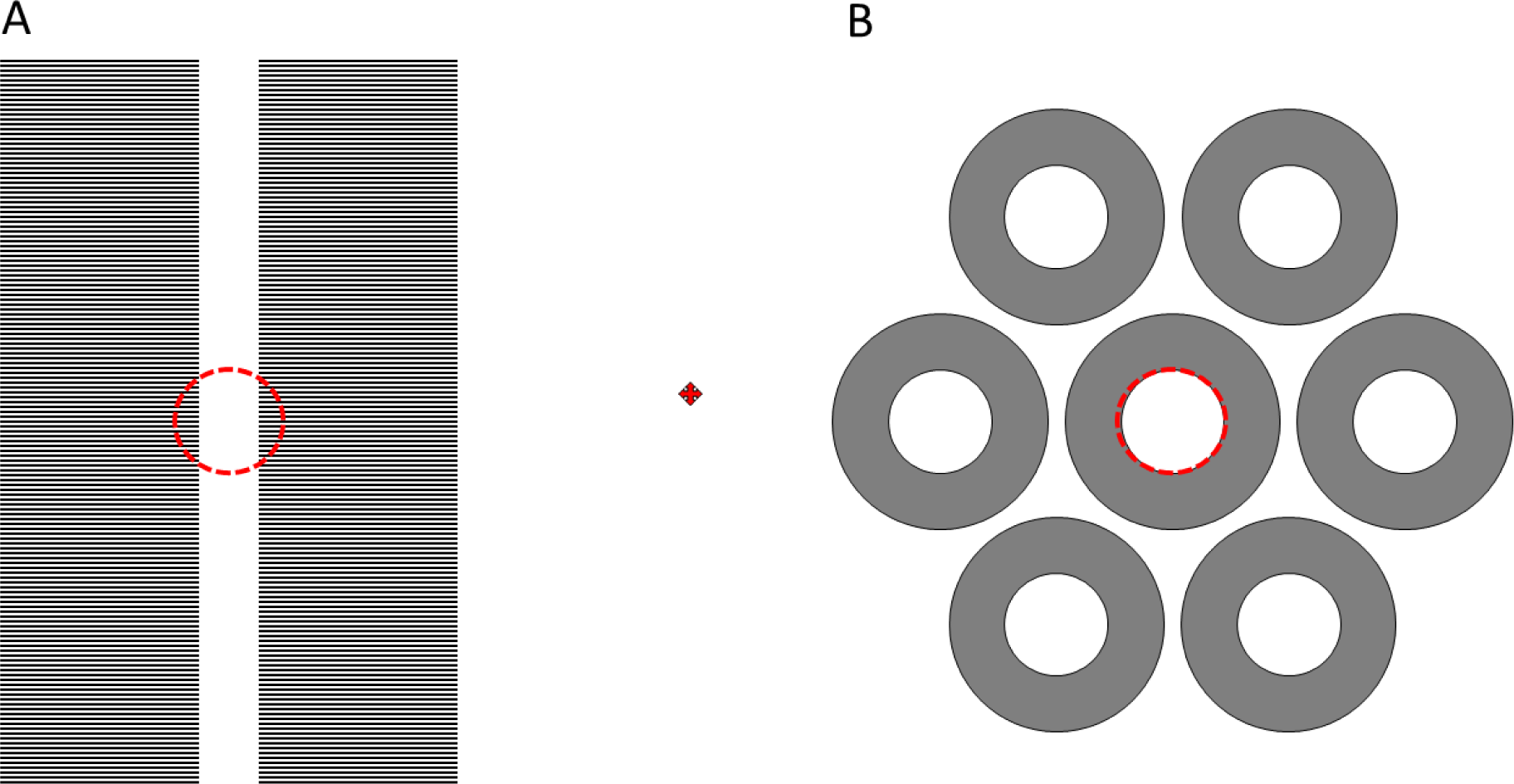
Filling-in at the blind spot operates at intermediate stages of visual processing (adapted from (Churchland & Ramachandran, 1996)). While fixating on the central red cross with one eye, adjust the page until the dashed red circles fall into the eye’s blind spot. **A.** Fixating with the *left* eye: The perceptual experience is of a continuous vertical white bar, second-order contours, rather than completing the horizontal grating texture. **B.** Fixating with the *right* eye: The central donut will appear perceptually completed as a filled grey disc, rather than a donut.

We note that some neuroimaging studies reported neural correlates of different types of filling-in in early visual cortex (Hong & Tong, 2017; Kok & de Lange, 2014; Weil & Rees, 2011). It therefore seems likely that multiple mechanisms can give rise to a filled percept, some of which operate actively at early stages of processing. Most relevant to our investigation, one study found a signature in early visual cortex of color filling of a shape outline (Hong & Tong, 2017). The stimulus design made it unlikely that these signals merely reflect differences in the inducing stimulus configuration. Therefore, the result implicates shape processing in producing the filled-in percept. However, given the higher-order nature of these shape stimuli, and the fact that similar results were observed across the visual hierarchy, suggests the V1 results could have involved feedback from higher regions. Moreover, this study used multivariate decoding techniques to evaluate the information about the color content of the filled percept. In contrast, we tested the presence of a signal that there is any stimulus visible within the blind spot at all. An equivalent decoding analysis for our research could entail decoding the direction of motion of our grating stimulus. It is possible that the V1 representation of the blind spot contains a signature of motion direction. Our experiment was not designed with such an analysis in mind; ideally one would present motion directions in longer trials to obtain robust signals.

Compelling evidence for the involvement of higher areas in perceptual filling-in of the blind spot is that adapting to a drifting grating annulus surrounding the blind spot, which creates a filled-in percept of a complete grating, produces a motion aftereffect at the blind spot location in the fellow eye (Murakami, 1995). Such inter-ocular transfer suggests the recruitment of binocular neurons, in higher extrastriate areas with large receptive fields covering both the blind spot and its surrounding space (e.g., MT). In fact, similar motion aftereffects can be induced at non-adapted locations that do not involve the blind spot (Snowden & Milne, 1997; Weisstein et al., 1977). It is therefore more parsimonious that filling-in occurs because these neurons merely discount the absence of signals from inside the blind spot and thus interpret the stimulus as a complete grating. This then leads to adaptation effects when stimulating the blind spot location.

Could the pattern of responses we observed be due to confounds in our design instead? First, we always stimulated the BE before the NE. This was necessary for delineating the blind spot border behaviorally and thus determine our stimulus dimensions, and for minimizing how often participants must be placed in the scanner bore. However, this could have introduced order effects, e.g., due to fatigue (Wylie et al., 2020). The NE eye was also deprived of vision for ∼45 minutes in the first half of each session. However, it is unlikely these effects are due to short-term monocular deprivation because this boosts the response of the deprived relative to the non-deprived eye (Binda et al., 2018).

Second, while the two eye conditions might have differed in terms of cognitive load or attentional allocation – in practice participants only needed to complete the filling-in task during BE viewing – they were monitoring for filling-in when viewing with either eye. It is implausible such attention would only affect higher-level regions but cause opposite effects in V1 and V2+3. While V4 and MT have been implicated with visual attention (O’Craven et al., 1999; Roe et al., 2012; Treue & Martinez Trujillo, 1999; Treue & Maunsell, 1999) these same attentional effects also affect lower visual areas; for instance, exogenous cuing affects V1 similarly to other downstream visual areas (Dugué et al., 2020). If responses in higher areas were driven by elevated visual attention in the BE condition, similar responses should also have been observed in pRFs outside the blind spot in V1_pRF_ and V2+3_pRF_. Instead, we found no differences or even subtly reduced responses there. Behavioral performance also indicated that attentional allocation and cognitive demand were comparable in both eyes (Supplementary Information). Crucially, the differential response in higher regions was greatest in the big condition, further inconsistent with an effect stemming from altered cognitive demands related to the filling-in task.

Third, researchers have noted a response bias in some animals of greater signals in the hemisphere contralateral to the stimulated eye (Schwarzkopf et al., 2007). However, such findings are specifically about early visual areas, and therefore cannot account for selective activity that excludes V1-V3. It also seems unlikely that this contralateral bias would manifest more strongly in higher cortical regions. Moreover, it is unclear if such a bias even exists in humans. It could be related to the degree of binocularity and the proportion of crossover fibers in the optic chiasm, which would predict it to be much less pronounced than in other species.

Finally, we also consider it unlikely that unstable eye fixation could explain our findings. Our experimental setup precluded eye tracking; however, participants performed a relatively demanding task at fixation throughout the main experiment, judging the direction of a small arrow target. Moreover, any break of fixation should have caused blurring of the stimulus-evoked responses in retinotopic cortex. In turn this might cause responses of pRFs estimated as falling into the blind spot, but we observed high retinotopic specificity of responses in the early visual cortex. In fact, the perceptual experience of a filled grating would also have been disrupted by any breaks in fixation.

While to our knowledge ours is the first brain imaging study to investigate the neural correlates of perceptual filling-in of the blind spot in human observers directly, a handful neuroimaging experiments have targeted the blind spot. When stimulating a large hemifield, there is no V1 signature of filling-in in fMRI responses, only a differential response indicating the blind spot representation (Tootell et al., 1998). Similarly, when the NE dominates perception during binocular rivalry, responses in the corresponding part of V1 are enhanced relative to when the BE is dominant; the V1 response therefore only reflects the retinal input of the currently perceived stimulus, not a filled-in percept (Tong & Engel, 2001). The cortical distance between activations evoked by stimuli on opposite sides of the blind spot also does not differ between blind spot and control eyes in either V1 or V2/3 (Awater et al., 2005). This shows there is no passive remapping of the blind spot where visual field maps on either side are “sewn-up” in this eye. That study, however, did not evoke filling-in. In general, these earlier fMRI studies lacked the spatial specificity possible today (Dumoulin & Wandell, 2008; Engel et al., 1997). This is crucial because the representation of the blind spot in human visual cortex is small, only subtending an area of about 50 mm^2^ (Adams et al., 2007). Moreover, no previous study has investigated regions beyond V2. We now reveal the brain activity measured during the actual percept of filling-in in unprecedented detail. Instead of any neural correlate of filling-in, we propose that neurons in higher extrastriate areas simply discount the absence of retinal inputs when integrating local signals. As Dennett argued (Dennett, 1992), it is the purpose of the visual system to find out what is out there in the world, rather than “filling-in” what is not there. Our finding is consistent with just such a mechanism whereby brain responses only reflect the available sensory evidence. Future research should employ experiments specifically designed to test this hypothesis further. Neural network models of perceptual processing could also prove instrumental for putting its predictions to the test.

## Data availability

Stimulus presentation functions and processed fMRI data and analysis code for reproducing the results in this study are publicly available at https://doi.org/10.17605/OSF.IO/DFMHN

National data protection rules prohibit the sharing of potentially identifiable brain images. Conditional sharing of raw data may be arranged with the corresponding author upon request.

## Acknowledgements

We thank Rasmus Pedersen for useful discussions of this work. This research was supported by an internal Faculty Research Development Fund from the *Faculty of Medical & Health Sciences* at the *University of Auckland* to DSS.

## Supplementary information

### Blind spot localization

Blind spot center eccentricity (in degrees of visual angle; mean; *M* = 16.74, Standard error of the mean; *SEM* = .36), and distance below the horizontal meridian (*M* = 1.34, *SEM* = .18), were consistent with known locations of the blind spot in visual space^30^. Mean blind spot height and widths in Session 1 (in degrees of visual angle - height: *M* = 6.59, *SEM* = .28; width; *M* = 5.63, *SEM* = .31), and Session 2, (in degrees of visual angle - height: *M* = 6.79, *SEM* = .25; width; *M* = 6.59, *SEM* = .28) also agreed with known dimensions of physiological scotomas. Mean area for the blind spot was 28.04 (*SEM* = 3.07) in Session 1, and 30.08 (*SEM* = 2.81) in Session 2. Visual inspection of Supplementary Figure S1 suggests there is variable agreement between sessions in the blind spot estimates, although the distance between the estimated centers of the blind spot from both sessions was always <1°. Importantly, the shape of the blind spot was generally consistent between sessions even if the exact location shifted. This is indicative of minor shifts in the field of view due to small variability in participants’ positions in the scanner between sessions rather than inaccuracies in determining the blind spot border. Note the plot for N02 in Supplementary Figure S1 only shows the blind spot from Session 1. This is because in the limited time in the scanner we were unable to use the Session 2 blind spot localization to effectively elicit filling-in but did find success using the Session 1 blind spot. Therefore, we used the blind spot for Session 1 in Session 2.

### Behavioral results

After collecting data from participant N97, we discovered that their responses were not recorded correctly due to a technical problem with the response box. Their data are still included in subsequent fMRI analyses because debriefing suggested they reliably perceived all stimulus conditions as intended. Furthermore, during debriefing participant N07 reported difficulty maintaining attention and complying with task instructions at several points during the scan, which was also reflected by their behavioral data. Thus, we performed subsequent analyses of the fMRI data once with all eight participants, and again after excluding participant N07. During viewing through the BE, the remaining participants reported seeing filling-in during an average of 88% (*SEM* = 5%) of the periods showing the filled stimulus, compared to 17% (*SEM* = 7%) and 10% (*SEM* = 5%) of big and small conditions, respectively. During viewing through the NE, participants reported seeing filling in during less than 1% of presentations across the three stimulus conditions. Filling-in was never reported during the blank period for either eye. We used JASP (Version 0.16.1) to further analyze arrow- and drift-subtasks with a two-way (eye; BE/NE, and stimulus condition; Blank/Big/Filled/Small) Bayesian repeated-measures analysis of variance (RM-ANOVA). Bayesian statistics allow statistical evidence in favor of the null hypothesis as well as direct comparisons between null and alternative hypotheses (Wagenmakers et al., 2018). Adopting Jeffreys’s criteria for interpreting Bayes factors (Jeffreys, 1961), we found moderate evidence for no effect of eye (BF_10_= .279), stimulus condition (BF_10_ =.241), and the interaction between eye and stimulus condition (BF_10_ =.018) in the arrow task.

### Results after excluding participant N07

After collating the behavioral results, we noted that participant N07 reported difficulty maintaining attention and complying with task instructions at several points during the scan, which was also reflected by their behavioral data. Hence, to investigate if their fMRI data affected the group analyses, we repeated those analyses after excluding N07. All results mirrored those of the original analysis in terms of the results of three RM-ANOVAs. For retinotopic areas V1_pRF_ and V2+3_pRF_ there was no positive βΔ for the filled condition. Results of the RM-ANOVA, for V1_pRF_ (Supplementary Figure S2A) showed a significant effect of location relative to blind spot boundary (F(1,6)=7.66, p=0.033), stimulus condition (F(2,12)=6.56, p=0.012), and the interaction term (F(2,12)=6.22, p=0.014). For V2+3_pRF_ (Supplementary Figure S2B) there was an effect of location relative to the blind spot boundary, (F(1,5)=13.01 p=.015), and stimulus condition (F(2,10)=10.9, p=.003), but no interaction (F(2,10)=0.17, p=0.849). Lastly, for the analysis incorporating all GLM-derived regions of interest (Supplementary Figure S2C), we also found an effect of region (F(4,24)=3.93, p=0.014), condition (F(2,12)=21.27, p<0.001), and the interaction term (F(2,12)=3.25, p=0.005). Lastly, conditions significantly different from zero (one-sample t-test vs. 0, false discovery rate-corrected p<.05) are labelled by asterisk in Supplementary Figure S2.

---

**Supplementary Table 1.** Selected visual areas as defined by Sereno et al. (2022) brain atlas. The left column shows areas pre-selected based on an initial identification of responsive areas, and the right column shows responsive clusters used as ROIs for analysis in the present study. Shaded regions across both columns denote the correspondence between pre-selected areas and labelled responsive areas consistently across participants. Note that V2 and V3 ROIs from the atlas were combined into the V2+3 cluster.

| Pre-selected Sereno areas | Cluster names |
| --- | --- |
| Early interparietal area |  |
| Anterior precuneus |  |
| Intraparietal sulcus |  |
| Frontal eye fields |  |
| Perifusiform gyrus |  |
| V4 | V4 |
| Superior temporal area |  |
| Parietal insular cortex |  |
| Aria prostratia |  |
| V6 |  |
| Middle temporal | MT |
| Lateral Occipital |  |
| Ventral occipital |  |
| V3A/B | V3A/B |
| V3 | V2+3 |
| V2 | V2+3 |
| V1 | V1 |

**Supplementary Figure S1.**
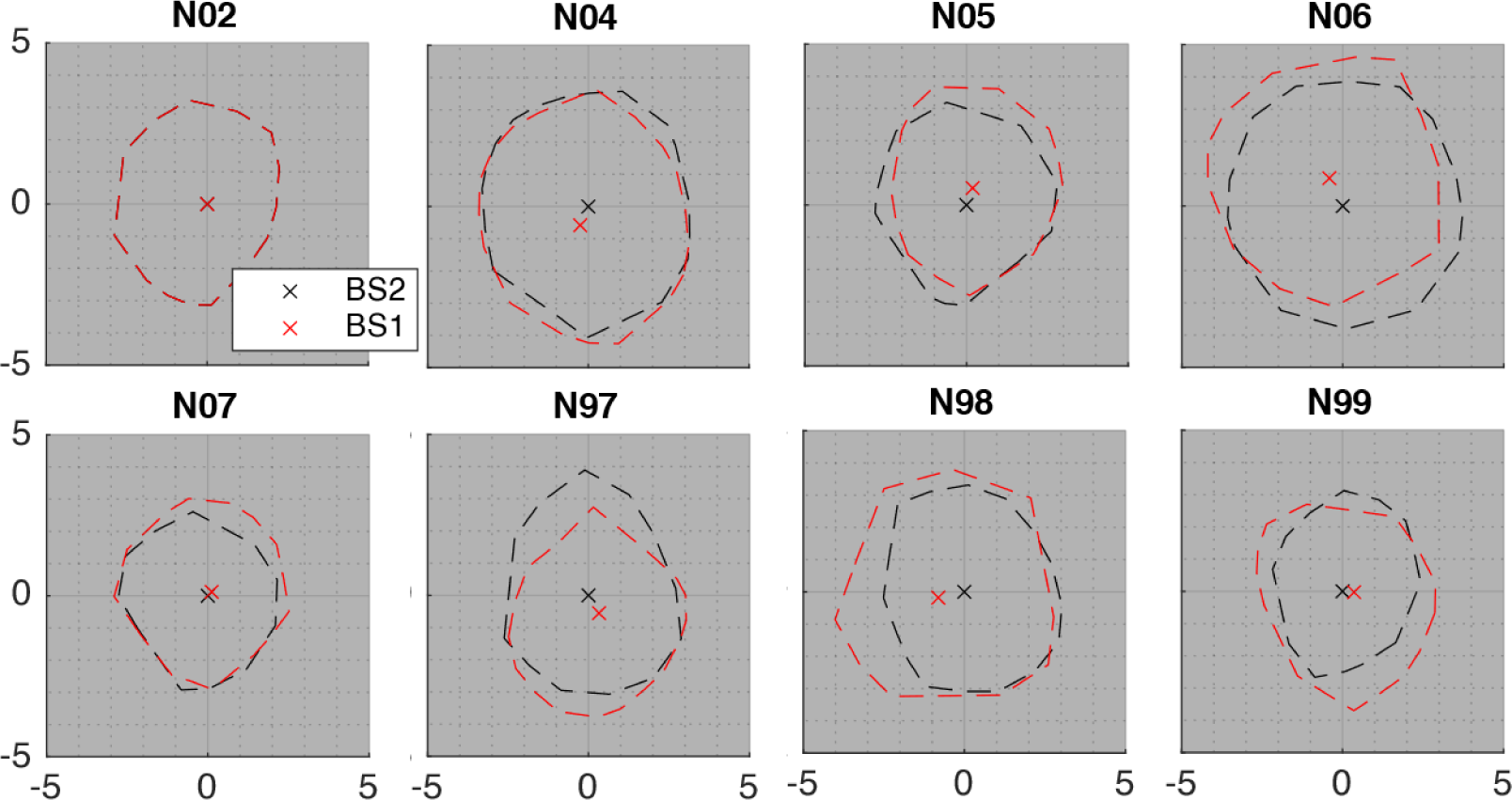
Blind spot plots for first (red: BS1) and second (black; BS2) sessions. Due to technical difficulties, for participant N1 we used the BS1 estimate for both sessions. Dashes show the boundaries and crosses the centers (calculated as the centroid) of the blind spot.

**Supplementary Figure S2.**
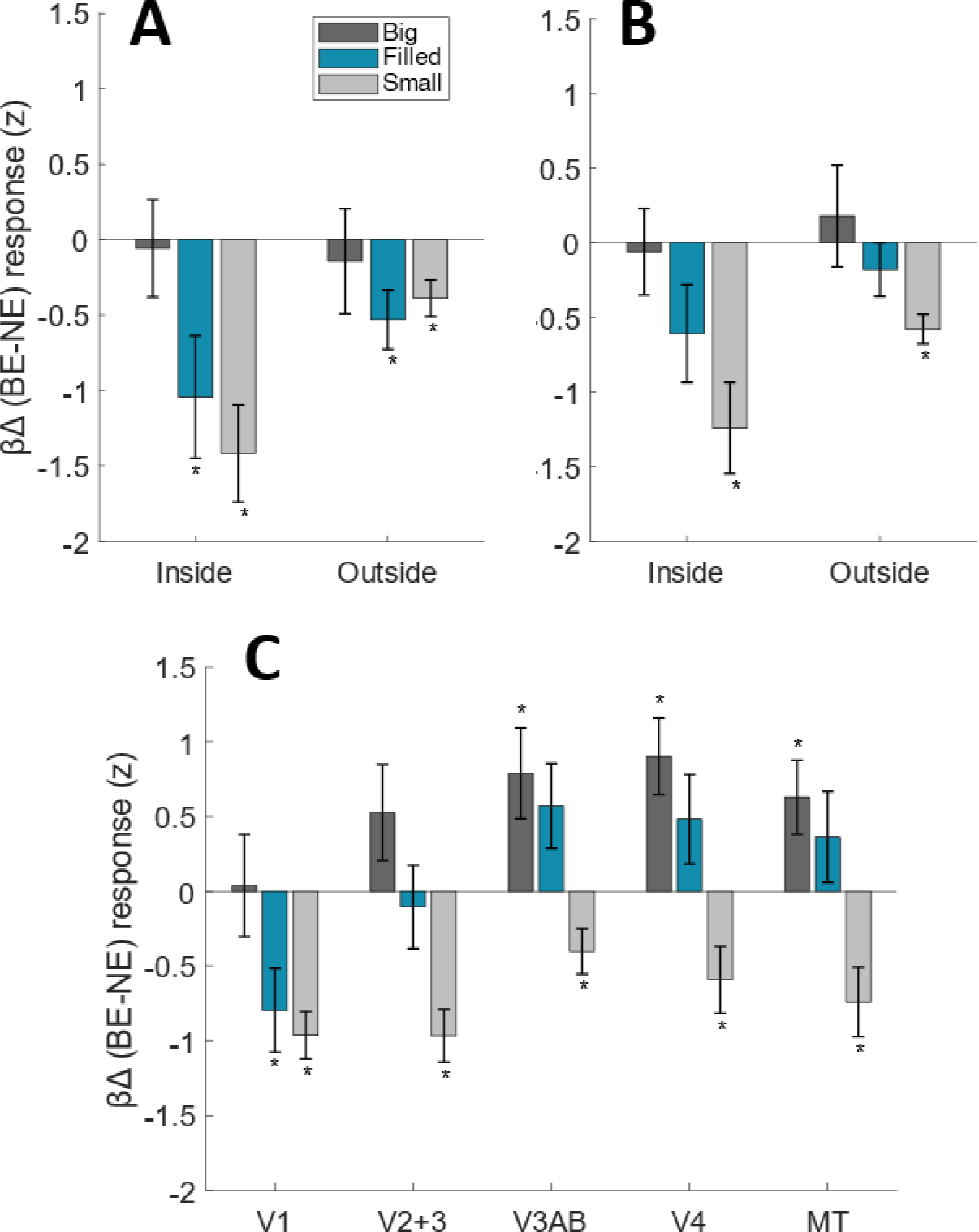
Difference in response to BE and NE stimulation (BE-NE βΔ) across regions of interest, excluding participant N07. Bars represent the average βΔ of all vertices within each region. **A.-B.** V1_pRF_ (**A**) and V2+3_pRF_ (**B**) data across stimulus conditions, inside and outside the blind spot. **C.** βΔ values in stimulus conditions for the five GLM-defined ROIs. Error bars represent ±1 standard error of the group mean. Bars labelled with a * differ significantly from zero (one-sample t-test against zero, false discovery rate-corrected *p* < .05).

